# Multivariate Transfer Entropy Quantifies Information Transfer Between Dyadic Partners During Real-Time Perceptual Decision-Making

**DOI:** 10.64898/2026.02.17.706300

**Authors:** J.P. Fiorenza, V Gnass, F Schneider, A Gail, F Wolf, S Treue, I Kagan, M. Wibral

## Abstract

Many social behaviours unfold continuously in time, yet most studies of social influence rely on discrete, trial-based paradigms. To quantify how humans actually integrate sensory and social signals during ongoing perceptual decision-making we here apply an information-theoretic analysis to data obtained with the Continuous Perceptual Report (CPR) paradigm. In the CPR task, pairs of participants tracked a dynamic random-dot motion stimulus, continuously reporting perceived direction and their confidence. Crucially, each participant had access to their partner’s moment-by-moment direction and confidence reports embedded within the stimulus display.

We quantified information flow between the stimulus and behavioural responses using multivariate transfer entropy, allowing us to detect whether participants acquired additional information from their partner once stimulus information had been received and integrated it into their behavioural response. We show that real-time social interactions predominantly occur within matching behavioural dimensions: partners’ perceptual choices influence participants’ own choices, and partners’ confidence influences participants’ confidence. Using both, more stimulus and social information contributed to improved task performance, and participants selectively acquired more information from better-performing partners.

Crucially, information integration was flexibly modulated by stimulus reliability: as sensory evidence became noisier, participants relied more on social information, but only when interacting with a highly reliable (e.g. computer-controlled) partner. Latency analyses further revealed that participants responded faster to changes in their partner’s perceptual choices of direction than to changes in stimulus direction. They also responded more slowly to changes in their partner’s confidence than to changes in their partner’s choice of direction. Together, these findings demonstrate that humans dynamically and adaptively integrate social information in a time-continuous manner, and they establish multivariate information-theoretic analyses as a powerful framework for studying real-time social cognition.

## 2 Introduction

In naturalistic conditions, social behaviour often unfolds in a time-continuous manner, allowing interacting individuals to dynamically modulate each other’s behaviour. Yet, such dynamic modulations have not been extensively studied until recently, as previous studies have mostly focused on trial-based, discrete paradigms. In these trial-based paradigms social context modulated perceptual reports and performance [1, 2, 3, 4, 5]. In particular, participants’ perceptual accuracy and confidence were modulated by the social setting – where the social settings were created by having pairs of participants, i.e. dyads, jointly perform a perceptual decision-making task.

More recently, experimental paradigms moved away from the discrete, trial-based structure, enabling the study of time-continuous perception through approaches such as continuous psychophysics [6, 7, 8]. Building on this line of work, Schneider and colleagues developed the Continuous Perceptual Report (CPR) task [5]. In a dyadic setting, human subjects continuously tracked the motion of a random dot pattern using a joystick with which they report their perceptual choice, via joystick direction, and their confidence, via joystick tilt, in real-time – while simultaneously observing their partners’ perceptual reports. Participants were incentivized to maximize their monetary reward score via peri-decision wagering, with their score dependent on a combination of accuracy and confidence. Importantly, there was no payoff interdependence between subjects.

By comparing subjects’ solo and dyadic performance, Schneider et al. observed that the presence of a co-acting social partner adaptively altered continuous-time perceptual decisions. Specifically, the worse-performing partner showed improvements in competence, confidence, and reward score, while the better-performing partner became less accurate and slightly less confident.

Here, we analyzed the CPR task using an information-theoretic approach that quantified information transfer between subjects’ behaviors with multivariate transfer entropy [9, 10, 11, 12]. Information theory has very successfully been applied in neuroscience (see [13] and references therein) and, more recently, to the study of naturalistic behaviour [14]. However, the use of multivariate transfer entropy to study the real-time behavioral integration of social influences remains unexplored. However, quantifying the effects of interacting partners on each other via measures of information flow indeed has been proposed as a first step toward describing the “interacting mind” [15].

In this work, we study information flow at the behavioural level in order to address four questions: (a) Do human subjects integrate, both, continuously changing stimulus- and partner-information into their behavioural readouts?, (b) Where does the information they integrate come from?, (c) How does the reliability of the stimulus modulate interaction between subjects?, (d) How fast do subjects integrate their partners’ behaviour?

We observed that real-time social information induces subjects to incorporate their partners’ perceptual decisions and confidence into their own corresponding behavioural dimension—that is, social influences occured within the same type of behavioural readout. Subjects integrated social information in a behaviourally relevant manner, improving their performance with increasing information transferred. Moreover, they flexibly adjusted to the reliability of the stimulus: the less reliable the stimulus was, the more they integrated information from their partners. Finally, we observed that subjects’ reactions to changes in their partners’ confidence occurred with longer lags than reactions to changes in their partners’ perceptual choices.

## 3 Methods and Materials

### 3.1 Study design and participants

Data were recorded from 38 human participants (Median age: 26.17 years, IQR 4.01; 13 of which with corrected vision) between January 2022 and August 2023. Prior to the experimental sessions, each participant was trained on two occasions for 1 hour. During the experimental phase, participants played three variations of the experimental paradigm: alone (solo), with another human player (human-human dyad), and (unknowingly to the participant) with a computer player impersonated by a human decoy (human-computer dyad). The experimental order was mixed and largely determined by the availability of participants. Most participants in this study originated from central Europe or South Asia. All procedures performed in this study were approved by the Ethics Board of the University of Göttingen (Application 171/292).

### 3.2 Experimental setup

Participants sat in separate experimental booths with identical hardware. They were instructed to rest their head on a chinrest, placed 57 cm away from the screen (Asus XG27AQ, 27” LCD). A single-stick joystick (adapted analog multifunctional joystick (Sasse), Resolution: 10-bit, 100 Hz) was anchored to an adjustable platform placed in front of the participants, at a height of 75 cm from the floor. The joystick was calibrated before data acquisition to ensure comparable readouts. Screens were calibrated to be isoluminant. Two speakers (Behringer MS16), one for each setup were used to deliver auditory feedback at 70 dB SPL.

The experimental paradigm was programmed in MWorks (Version 0.10 – https://mworks.github.io). Two iMac Pro computers (Apple, MacOS Mojave 10.14.6) served as independent servers for each setup booth. These computers were controlled by an iMac Pro (Apple, MacOS Mojave 10.14.6). Custom-made plugins for MWorks were used to generate and display the stimuli, to handle the data acquisition from the joystick (100 Hz sampling rate), and to incorporate all data from both servers into a single data file.

### 3.3 Continuous perceptual report (CPR) game

Participants were instructed to maximize monetary outcome in a motion tracking game. In this game, subjects watched a frequently changing random dot pattern on the screen and used a joystick to indicate their current motion direction perception. The joystick controlled an arc-shaped response cursor on the screen (partial circle with fixed eccentricity, Solo: 2 degree of visual angle (‘dva’) radius from the center of the screen; Dyadic: 1.8 dva and 2 dva radius). The angular direction of the joystick was linked to the cursor’s polar center position. In addition, the joystick tilt was permanently coupled to the cursor’s width (see below, 13 – 180 degrees). This resulted a continuous representation of the joystick position along its two axes. By moving the joystick, participants could rotate and shape the cursor. At unpredictable times (1 percent probability every 10 ms), a small white disc (‘reward target’, diameter: 0.5 dva) appeared on the screen for a duration of 50 ms at 2.5 dva eccentricity, congruently with the motion direction of the stimulus. Whenever a target appeared in line with the cursor, the target was considered collected (‘hit’) and the score of the participant increased. To help participants performing such alignment to the best of their perceptual abilities, a small triangular reference point was added to the center point of the cursor. Throughout the experiment, participants were required to maintain gaze fixation on a central fixation cross (2.5 dva radius tolerance window) or the cursor would disappear and no targets could be collected until fixation was resumed.

In the solo experiments, the cursor was always red. In dyadic conditions, the two cursors, present on screen simultaneously, were red and green (isoluminant at 17.5 cd/m2). During dyadic experiments, the position of the two cursors switched between stimulus cycles, with the red cursor always starting on top (i.e. more eccentric), but not overlapping the green cursor. Each cursor color was permanently associated with one of the two experimental booths. After the mid-session break, participants switched booths, contributing an equal amount of data for each setup (600 reward targets, 20 min, up to 17 stimulus cycles). Players initiated new stimulus cycles with a joystick movement. Each stimulus cycle could last up to 75 seconds, during which the RDP’s motion direction and coherence changed at pseudorandomized intervals, resulting in the presentation of 30 stimulus states per cycle.

### 3.4 Random dot pattern (RDP)

We used a circular RDP (8 dva radius) with white dots on a black background. Each dot had a diameter of 0.1 dva, moved with 8 dva/s and had a lifetime of 25 frames (208 ms). The overall dot density was 2.5 dots/dva^2^. The stimulus patch was centrally located on the screen, and its central part (5 dva diameter) was blacked out. In this area we presented the fixation cross and the response arc. The RDP motion direction was randomly seeded and set to change instantly by either 15 deg, 45 deg, 90 deg or 135 deg after a pseudo-randomized time interval of 1250 ms to 2500 ms, with equal probabilities for clockwise or counterclockwise direction changes. These changes only affected the dots carrying the motion signal. The dots’ coherence changed pseudo-randomly after 10 RDP direction changes to the coherence level that was presented least. Seven coherence conditions were tested: 0, 8, 13, 22, 36, 59, 98%. For analysis, we calculated the instantaneous (frame-to-frame) direction and coherence of the random dot pattern by integrating the individual movements of dots in the stimulus patch into a resultant vector. Dots that moved over the stimulus boundaries or expired due to the dot lifetime between frames were excluded. The resultant vector direction and length summarize the motion energy between two frames.

### 3.5 Gaze control

Participants were required to maintain gaze fixation at the center of the screen throughout each stimulus cycle. We used a white cross (0.3 dva diameter) as anchor point for the participants’ gaze. The diameter of the fixation window was set to 5 dva. An eye tracker (SR research, EyeLink 1000 Plus) was used to control gaze position in real-time. If the gaze position left the fixation window for more than 300 ms, the player’s arc would disappear from the screen, preventing target collection. In addition to this, an increase of the fixation cross’ size, together with a change in color (white to red), signaled to the participants that fixation was broken. As soon as the gaze entered the fixation window again, visual parameters were reset to the original values and the arc would reappear, allowing the player to continue target collection.

### 3.6 Reward score

Participants were incentivized to collect as many targets as possible with the highest possible score by coupling the sum of score to a proportional monetary reward.

The target score was computed from the minimum polar distance between the arc’s center position and the nominal direction (target center, ‘accuracy’) as well as the angular width of the response cursor (‘joystick tilt’) at the moment of collection as:

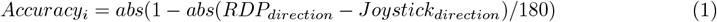

where *RDP*_*direction*_ refers to the direction of the random dot pattern and Joystick refers to the direction of the joystick at sample i.

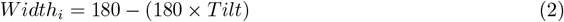

where tilt varies between 0 and 1 for minimal and maximal radial joystick positions, respectively.

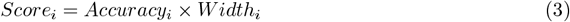

Thus, narrower and more accurately placed cursors caused higher reward scores.

### 3.7 Feedback signals

Various feedback signals were provided throughout the experiment to inform participants about their short- and long-term performance. All feedback signals were mutually visible.

#### Immediate feedback

Immediately after a target was collected, visual and acoustic signals were provided simultaneously. The auditory feedback consisted of a 200 ms long sinusoidal pure tone at a frequency determined by the score. Each tone corresponded to a reward range of 12.5 percent, with lower pitch corresponding to low reward score. We used 8 notes from the C5 major scale (523, 587, 659, 698, 784, 880, 988, 1047 Hz). Sounds were on- and off-ramped using a 50 ms Hanning window. No sound feedback was given for missed targets. In solo experiments, the visual feedback consisted of a 2 dva wide circle, filled in proportion to the score with the same color as the arc’s player. The circle was presented in the center of the screen, behind the fixation cross for 150 ms. In dyadic conditions, the visual feedback consisted of half a disc for each player and both color-coded halves were mutually visible.

#### Short-term feedback

During each stimulus cycle, a running average of the reward score was displayed for each player with a 0.9 dva wide, color-coded ring around the circumference of the RDP (18.2 dva and 19.4 dva diameter respectively for each player). After every target presentation, the filled portion of the ring updated. To avoid spatial biasing, the polar zero position of the ring changed randomly with every stimulus cycle.

#### Long-term feedback

Cumulative visual feedback was provided after each stimulus cycle (during the inter-cycle intervals) for 2000 ms. It displayed the total reward score accumulated across all cycles as a colored bar graph located at the center of the screen. A grey bar (2 dva wide, max height: 10 dva) indicated the maximal possible cumulative score after each cycle. A colored bar next to it (same dimensions) showed how much was collected by the player so far. In dyadic experiments, red and green bars would be shown on either side of the grey bar. In solo experiments, a red bar was shown to the left of the grey bar. The configuration of the visual stimuli and task parameters is illustrated in Figure 1 and as a supplementary video file Supplementary Video 1.

**Figure 1:**
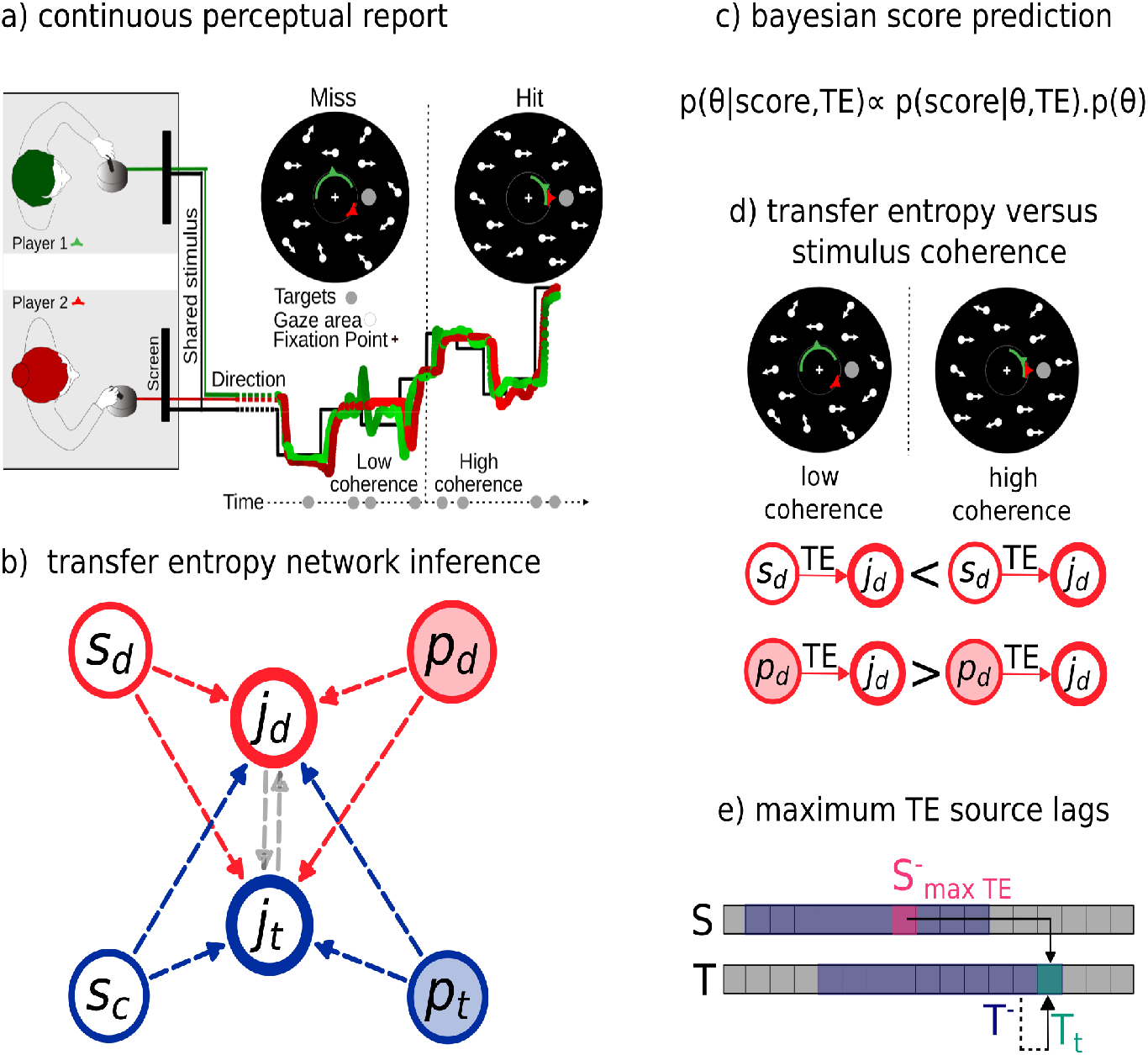
Analysis workflow. **(a)** Continuous Perceptual Report task (adapted from Figure 1 in Schneider et al 2024). Two participants sat in adjacent experimental booths and played a motion tracking game either alone (‘solo’) or together with a partner (‘dyadic’, mixed order). They used a joystick to track a random dot pattern for monetary reward (referred as joystick direction, *j*_*d*_). The random dot pattern (‘RDP’) was presented with a circular aperture and with a blacked out central fixation area. Subjects were instructed to look at the central fixation cross. The motion stimulus was continuously presented in intervals of about 1 minute and changed frequently in motion direction and coherence level. In dyadic experiments, the visual stimulus was shared. Joystick-controlled cursors of both players as well as visual feedback were mutually visible in real-time. The joystick tilt (*j*_*t*_) was linked to size of the cursor: less tilt – wide; more tilt - narrow. The color-coded cursors were located at the edge of the fixation area. The stimulus motion signal was predictive of the location of behaviorally relevant reward targets (gray disc). Alignment of the cursor arc with the target resulted in target collection (‘hit’) and reward. Target presentation occurred in pseudorandom time intervals. The visual and auditory feedback representing the reward score was provided after each target hit. **(b)** Multivariate transfer entropy (TE) was used to identify which of the information sources – stimulus direction (*s*_*d*_), stimulus coherence (*s*_*c*_), partner’s direction (*p*_*d*_) and partner’s tilt (*p*_*t*_) – transferred information to the subject joystick’s response – direction (*j*_*d*_) and tilt (*j*_*t*_). **(c)** Bayesian linear regression model was used to estimate whether the information transferred – i.e., transfer entropy – predicts subject’s performance. **(d)** Transfer entropy modulation by stimulus coherence was studied for two connections: (1) stimulus direction to joystick direction (*sd* →*jd*) and (2) partner’s direction to joystick direction (*pd*→ *jd*). We hypothesize that TE will increase in *sd* →*jd* with increasing stimulus coherence, and decrease in *pd* →*jd* for decreasing stimulus coherence. **(e)** Lags with maximum information transferred were extracted from each detected connection.

### 3.8 Transfer Entropy Network Identification

In this study we analyzed the information transfer between various variables describing the stimulus as well as human behaviour. In our context specifically, the presence of significant information transfer means that some information available in a source signal, i.e. the stimulus or a partner’s behaviour, has an influence on a (behavioral) target variable. Thus, the difference between presence and absence of information transfer lies in the target that adapts to the incoming information (or doesn’t) rather than in the presence or absence of information in the source.

In terms of algorithms, we used a multivariate form of transfer entropy (TE) as a measure of information transfer to detect which of the stimulus dimensions (stimulus direction and stimulus coherence) and behavioural responses (subject’s direction or subject’s tilt) transmitted information to each behavioural response. The multivariate approach [11, 12] allows us to detect information sources (stimulus dimensions and behavioural responses) in the context of the others, thus avoiding double counting of information redundantly present in multiple sources and revealing potential synergies between information sources. Most importantly for our study, we can thus identify information flowing between subjects (e.g., from the partner’s direction to a subject’s direction) while fully accounting for the information they have already received from the stimulus.

Transfer entropy (Eq 4) is a lag-based measure derived from mutual information that requires both a suitable embedding to reconstruct the state space from a set of observations and an estimation of probability densities of these states.

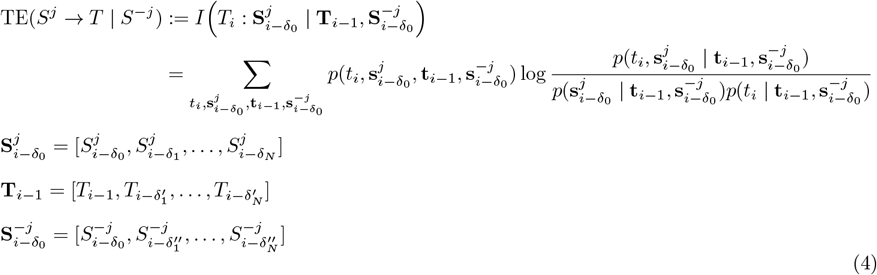

where *i* indexes the discrete time steps, 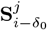 stands for the state vector of source process *j* with its past state variables indexed with *δ* starting at *δ*_0_, **T**_*i*−1_ the state vector of the target with its past state variables indexed with *δ*^′^ starting at 1 time step in the past, 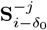 stands for the state vector of the rest of the source processes – i.e., all except *j* – with its past state variables indexed with *δ*^″^ starting at the same *δ*_0_ as the source process *j*. Lowercase letters represent the realizations of these random variables 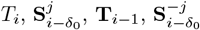.. Bold case letters indicate vector notation.

As seen in Eq. 4, transfer entropy quantifies how much information the past of a source *S* ^*j*^ contains about the present of a target T, given the target’s past and the rest of the sources (indicated with *-j*). Therefore, quantifying information in the past of *S* ^*j*^ not redudantly contained in the target’s past nor in the past of the rest of the sources *S*^−*j*^. Thus, a significant TE indicates the presence of information flowing from *S* ^*j*^ to T in the context of *S*^−*j*^.

In this work, we used the Gaussian TE estimator, [16, 17]. Lags (delayed versions of the original signal, represented in Eq. 4 by using *δ*_*l*_, 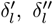 with l=1…N) were selected using a non-uniform embedding approach. The non-uniform embedding approach consists of selecting those lags – i.e., *δ*_*l*_ – which are maximally informative in the context of already identified lags. Each new lag is included iteratively following a greedy forward-selection strategy that maximizes conditional mutual information. Maximally informative lags for the sources were searched within a window of 200 to 1300 ms in the past, and for the target within a window of 1 to 1000 ms in the past. The greedy algorithm as well as the hierarchical statistical testing scheme that handles the family-wise error rate is explained in more detail in [11].

After detecting the source-target links across all dyads, we classified the dyads based on their social interaction. If both subjects influenced each other (i.e., links were present in both directions), the dyad was classified as a mutual influence dyad. If the influence was only from one player to the other, the sender was classified as leader and the receiver as follower. The classification was carried out separately for the subjects’ direction and subjects’ tilt.

All the information-theoretic analyses were run using the toolbox IDTxl [12] which internally makes use of the Gaussian estimator implemented in JIDT [18].

### 3.9 Bayesian linear models of subjects’ performance

From the above transfer entropy (TE) analysis we identified the information sources (i.e., which stimulus and behavioural dimensions transferred information) and quantified the amount of information they transferred to their respective targets. We then used Bayesian linear regression models to investigate whether, and to what extent, subjects made use of this information to improve task performance. Specifically, we first modeled subjects’ scores as a linear combination of TE values originating from different stimulus and social sources. Additionally, we examined whether the amount of information subjects received from their partner’s direction depended on the partner’s relative solo performance. The following sections describe these models and the estimation procedure.

#### 3.9.1 Models predicting the subject’s performance via transfer entropy

We implemented four linear regression models with parameters estimated using Bayesian inference with Markov chain Monte Carlo (MCMC) sampling. In all four models, we used a Gaussian likelihood (Eq 5) to describe the data given the parameters, and we modelled the means, *µ*, of the Gaussian likelihood distributions as a linear combination of TE from different stimulus and social information sources. First, we used only information transfers originating from the stimulus as predictors of *µ* (SI model), i.e., the TE from stimulus direction to joystick direction (*TE*_*sd*→*jd*_) and from stimulus coherence to joystick tilt (*TE*_*sc*→*jt*_) (Eq 6). Second, we used information transfers from both the stimulus and social information sources as predictors (SSI model), i.e., the TE from four links: (a) stimulus direction to joystick direction, (b) stimulus coherence to joystick tilt, (c) partner’s direction to joystick direction (*TE*_*pd*→*jd*_), (d) partners’ tilt to player’s tilt (*TE*_*pt*→*jt*_) (Eq 7). Third, we used the same predictors as in the second model but implemented them hierarchically (HSSI model) based on the type of dyad (i.e., human-human or human-computer dyad, see section 3.2) for the partner’s direction to joystick direction (Eq 8). In this hierarchical formulation, the effect of *TE*_*pd*→*jd*_ was modelled separately for the two dyad types, with distinct regression coefficients for human-human 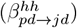 and human-computer 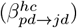 dyads. Our final model was a null model including only the intercept *β*_0_ as a predictor.

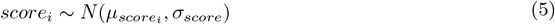

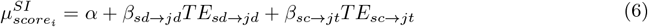

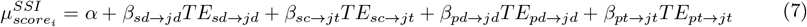

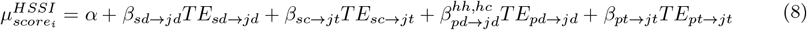

#### 3.9.2 Model predicting transfer entropy via relative solo performance

We used a Bayesian linear regression to predict the transfer entropy from the partner’s direction to the subject’s joystick direction based on the relative solo performance of the dyad members.

Transfer entropy values were modelled with a Gaussian likelihood (Eq. 9), where the mean *µ* was expressed as a function (Eq. 10) of the difference between the partner’s solo score (*S*_*p*_) and the subject’s solo score (*S*_*j*_). Note that *S*_*j*_ represents the solo score of the subject who receives the partner’s information.

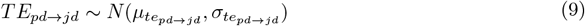

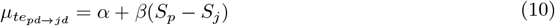

#### 3.9.3 Estimation of model parameters, priors and model comparison

Our Bayesian inference analysis was performed using the Python package PyMC [19] with NUTS (No-U-Turn Sampling) [20] and multiple independent Markov chains. We ran 10 chains, each performing 1000 burn-in (tuning) steps to allow the chains to equilibrate, followed by 10000 steps to sample from the approximate posterior distribution.

Before sampling, players’ scores and transfer entropy estimates were standardized. Priors were centered at zero to encode symmetry between positive and negative effects, reflecting no prior preference for increases or decreases in subjects’ scores. The priors were chosen to be weakly informative relative to the likelihood, remaining sufficiently diffuse to allow the data to dominate posterior inference. Prior predictive checks confirmed that the priors generate a broad range of outcome values (Fig. S2a). Priors can be found in Table 1.

**Table 1:** Prior distributions for all model parameters: score prediction models (SI, SSI, HSSI) using transfer entropy, and the model predicting transfer entropy from relative solo performance.

| Parameter | Prior distribution |
| --- | --- |
| $\sigma_{score}$ | HalfCauchy($\beta=2$) |
| $\sigma_{te_{pd \rightarrow jd}}$ | HalfCauchy($\beta=2$) |
| $\alpha$ | Normal($\mu=0, \sigma=4$) |
| $\beta_{sd\_jd}$ | Normal($\mu=0, \sigma=4$) |
| $\beta_{sc\_jt}$ | Normal($\mu=0, \sigma=4$) |
| $\beta_{pd\_jd}$ | Normal($\mu=0, \sigma=4$) |
| $\beta_{pt\_jt}$ | Normal($\mu=0, \sigma=4$) |
| $\beta$ | Normal($\mu=0, \sigma=4$) |

We evaluated the model’s performance using the leave-one-out (LOO) cross-validation criteria [21]. This approach compares models by their out-of-sample predictive accuracy, i.e., their log point-wise predictive density (lpd)

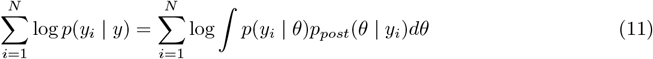

with N being the total number of data points, *y*_*i*_ a single data point, *p*(*y*_*i*_ | *θ*) its likelihood and *p*(*θ* | *y*_*i*_) the posterior over the parameters *θ*.

The pointwise out-of-sample prediction accuracy was approximated using LOO, which computes (Eq 11) individually for each left-out data point based on the model fit to the other data points.

### 3.10 Effects of modulation of stimulus coherence on information transfer

We studied the effects of the modulation of stimulus coherence on the TE of two links: from stimulus direction to joystick direction, and from partner’s direction to joystick direction. Transfer entropy was estimated using the embedding previously inferred in the network analysis (section 4.1). TE values were then recomputed separately for each of the seven coherence levels (see section 3.4). The analysis included all human participants in both human–human and human–computer dyads.

We estimated the mean difference between TE at each coherence level using Bayesian regression. TE at each coherence level was modeled with a Gaussian likelihood, where the mean, *µ* (Eq 12), was described as the sum of a grand mean, *µ*, and a deviation, *α*_*i*_, from it. For a pair of coherence levels, e.g., 0.08 and 0.13, we computed the difference in TE as *α*_0.08_ - *α*_0.13_.

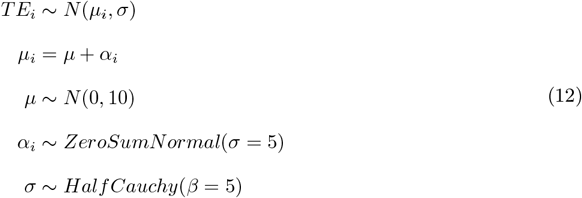

### 3.11 Maximum information transfer lags

From the set of source lags transmitting information to the target, we selected those corresponding to maximum TE. This was done for the four main links detected: (a) sd → jd, (b) sc → jt, (c) pd → jd and (d) pt → jt. We then used Bayesian regression to estimate the mean difference between the lags with maximum information transfer, e.g., sd → jd vs pd → jd. We modelled the lags using a Gaussian likelihood, with its mean, *µ* (Eq 13), described as the sum of a grand mean, *µ*, and a deviation *α*_*i*_, from it. For a a pair of links, e.g., sd → jd vs pd → jd, we computed the difference in lags as *α*_*sd*→*jd*_ −*α*_*pd*→*jd*_.

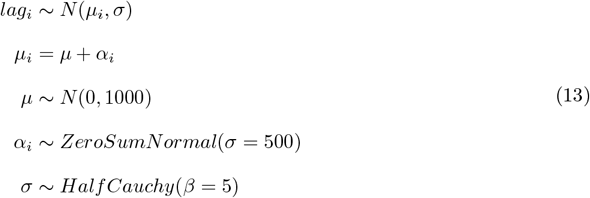

## 4 Results

### 4.1 The structure of information flow from stimulus dimensions to and between behavioral dimensions

Social information was predominantly transferred between players in matching behavioral dimensions, i.e., from joystick direction to joystick direction (p < 0.05, FDR-corrected, 90% of subjects), and from joystick tilt to joystick tilt (p < 0.05, FDR-corrected, 38% of subjects) in real human-human dyads. In contrast, between non-matching behavioral dimensions, such as from one player’s joystick direction to the other’s joystick tilt, less instances of significant information transfer were observed; for example, in only 4% of the subjects their joystick tilt received information from their partner’s joystick direction (Fig. 2a).

**Figure 2:**
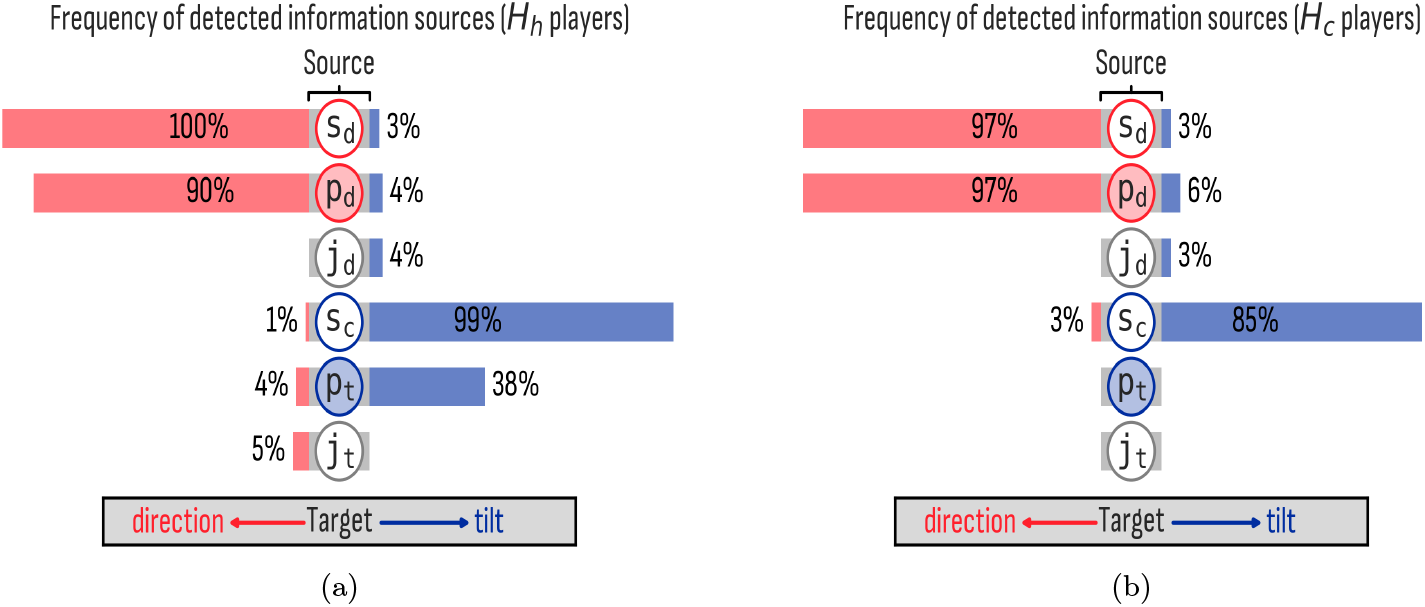
Frequency of detected influences across subjects in human-human (*H*_*h*_) and human-computer (*H*_*c*_) dyads. (a) Human-human dyads, (b) human-computer dyads. Percentage of subjects’ joystick direction and joystick tilt with a significant transfer entropy (p<0.05, FDR-corrected) from each candidate source: stimulus direction (*s*_*d*_), partner’s direction (*p*_*d*_), joystick direction (*j*_*d*_), stimulus coherence (*s*_*c*_), partner’s tilt (*p*_*t*_), joystick tilt (*j*_*t*_). Joystick direction and tilt refer to the subject’s behavioural dimension. For example, only 4% of subjects’ joystick direction sent information to its own joystick tilt in human-human dyads.

Both, in human-human (*H*_*h*_) dyads and in human-computer (*H*_*c*_) dyads where subjects (unknowingly) played together with a simulated, highly accurate partner who exhibited a constant, intermediate level of confidence, we observed significant transfer entropy (p < 0.05, FDR-corrected) from stimulus direction to joystick direction in almost all human players (100% in *H*_*h*_ and 97% in *H*_*c*_ dyads).

However, as required on theoretical grounds, there was a difference in terms of the information flowing to subjects’ joystick tilt between human-human dyads and in human-computer dyads. As the computer player in an *H*_*c*_ dyad did not adapt its “confidence” to the stimulus coherence, and hence does not provide information, none of the human subjects received information from the computer’s joystick tilt in the *H*_*c*_ dyads. In addition, the constant computer-generated joystick tilt was an isolated process in almost all dyads; i.e., it did not receive information from nor send information to any process, whether stimulus or social (Fig 2b). These two results suggest that the information-theoretic analysis indeed has a low false-positive rate.

Furthermore, when playing with a simulated partner, there was a slight decrease in the number of players (85%) whose joystick tilt received information from stimulus coherence (Fig. 2b), suggesting that some players were influenced by the absence of confidence signaling from the presumed partner.

Quantifying interactions in terms of information transfer using transfer entropy allowed us to identify the information sources influencing each behavioural dimension, as well as to detect distinct interaction patterns across dyads. Based on these patterns, we classified dyads into three categories for each behavioural dimension (i.e., joystick direction and tilt): (a) *mutual influence*, when both players send and receive information from each other; (b) *leader–follower*, when only one player sends information (the leader) and the other only receives it (the follower); and (c) *no interaction*, when no information flows between players.

Mutual influence via joystick direction was present in 81.2% of dyads, via joystick tilt in 20.8% and in both dimensions in 17%. Leader-follower dynamics were observed via joystick direction in 8.3%, via players’ joystick tilt in 16.7% and in both dimensions in 4%. Only 2.1% showed no interaction in joystick direction, whereas 45.8% showed no interaction via joystick tilt, indicating a predominant interaction via players’ joystick direction compared to tilt (Fig. 3).

**Figure 3:**
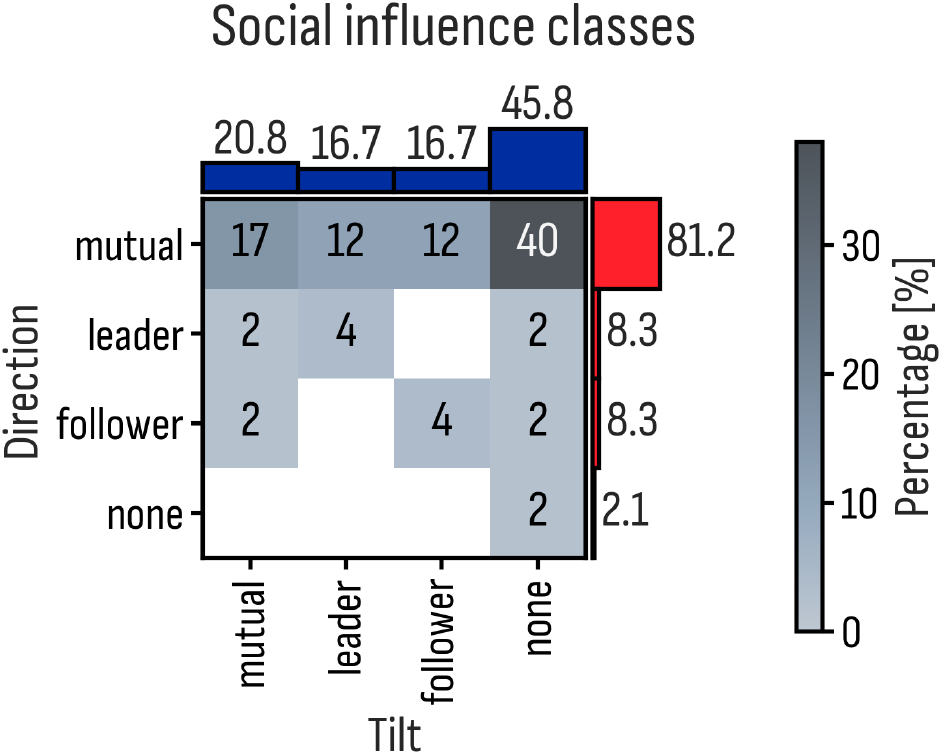
Social classification of human-human dyads based on information transfer between subjects’ joystick direction and joystick tilt. Social influence classes were mutual influence (mutual), leader-follower or none (see section 3.8). In the x-axis percentage of subjects on each category based on their joystick tilt (e.g., 16.7% leader-follower). In the y-axis percentage of subjects based on their joystick direction (e.g., 8.3% leader-follower). Values within the matrix represent the joint classification based on direction and tilt (e.g., 17% of subjects were belonged to mutual influence both in direction and tilt).

High joystick accuracy correlated with high tilt across all levels of task difficulty (Supplementary Figure 2 in Schneider et al 2025). This justifies the assumption that joystick tilt can be interpreted as proxy measure for perceptual confidence, as its placement seems to be guided by a peri-decision wagering via a meta-cognitive assessment of real-time perception. Therefore, our results can be interpreted such that information about a partner’s confidence is integrated in a subject’s own confidence-assessment in the real-time social interactions of our CPR task.

### 4.2 Information transfer improves player’s score

To determine whether stimulus (direction, coherence) and social (direction, tilt) information influence subject’s performance, we predicted their reward score (i.e., performance in the the dyadic setting) as a function of the transfer entropies from the stimulus and social information sources (see section 3.9 for the models’ description).

We observed an increase in subjects’ score with greater transfer entropy from stimulus direction to joystick direction (*β*_*sd*_*jd*_ = 0.60, [0.42, 0.77] 94% credible interval), from the partner’s direction to joystick direction (*β*_*pd*_*jd*_ = 0.63, [0.49, 0.78] 94% credible interval), and from stimulus coherence to joystick tilt (*β*_*sc*_*jt*_ = 0.17, [0.005, 0.342] 94% credible interval), as shown by the SSI model (Fig. 4). When accounting for partner type (human or computer) by modelling the coefficient *β*_*pd*_*jd*_ hierarchically across dyad types, the same increase in subjects’ scores with increasing TE was observed.

**Figure 4:**
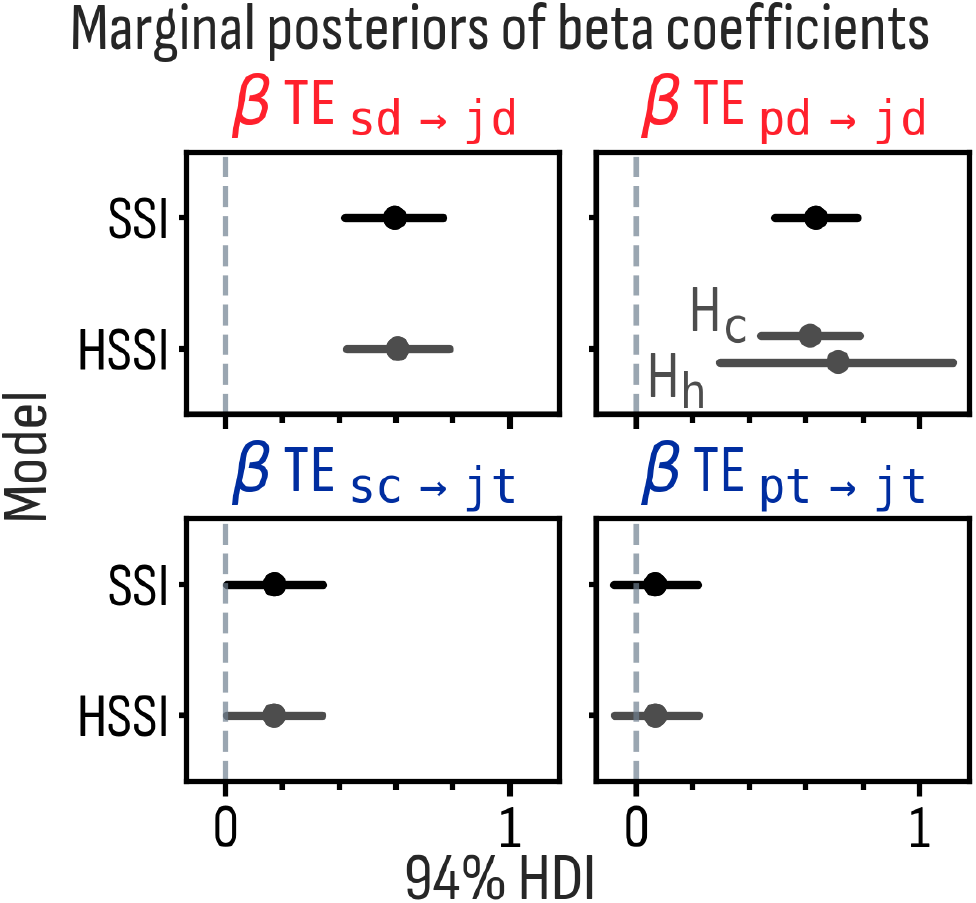
Posterior distribution of parameters in both stimulus-social model (SSI) and hierarchical, stimulus-social model (HSSI). 94% highest density interval (HDI) shows a positive effect indicating an increase in subjects’ score with increasing transfer entropy from stimulus direction 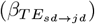, partner’s direction 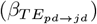 and stimulus coherence 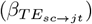.

Models incorporating both social and stimulus information (i.e., SSI and HSSI models) predicted players’ scores better than models accounting for stimulus information alone (SI), as indicated by the larger expected log pointwise predictive density (ELPD) for SSI and HSSI compared to SI (Fig. S1 in supporting information).

The fact that the SSI model performed slightly better than the HSSI model suggests that the evidence in our data was not enough to outweigh model complexity, resulting in a better performance for the simpler model (SSI, non-hierarchical).

The observed data was informative about all four parameters (*β*_*sd*→*jd*_, *β*_*sc*→*jt*_, *β*_*pd*→*jd*_, *β*_*pt*→*jt*_), as indicated by the difference between priors and posteriors (Fig. S2 c,d). Besides, both SSI and HSSI models were able to generate reward score values compatible with those contained in the data; as observed by the overlap between the data and scores generated after sampling from the posterior distributions (Fig. S2 b).

Taken together, these results suggests that subjects integrate both stimulus and social information, leading to improved task performance as the amount of information received increases.

#### 4.2.1 Information acquired from partners increases with their performance

Since greater acquisition of social information from a partner’s joystick direction 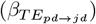 was associated with higher subject scores (see Fig. 4), we next asked whether the amount of information subjects acquired from their partner depended on the partner’s relative performance. Previous work has shown that social modulation (manifested as larger discrepancies between subjects’ accuracy and confidence in solo versus dyadic settings) increases with greater differences in partners’ solo performance [5].

Consistent with these findings, we observed that subjects paired with better-performing partners (indicated by positive values of “Δ *solo reward score*” in Fig. 5) received more information from their partner’s joystick direction. Specifically, transfer entropy from partner direction to joystick direction increased linearly (*β* = 0.44, 94% HDI [0.25, 0.63]) as a function of the difference in solo performance between members of a dyad.

**Figure 5:**
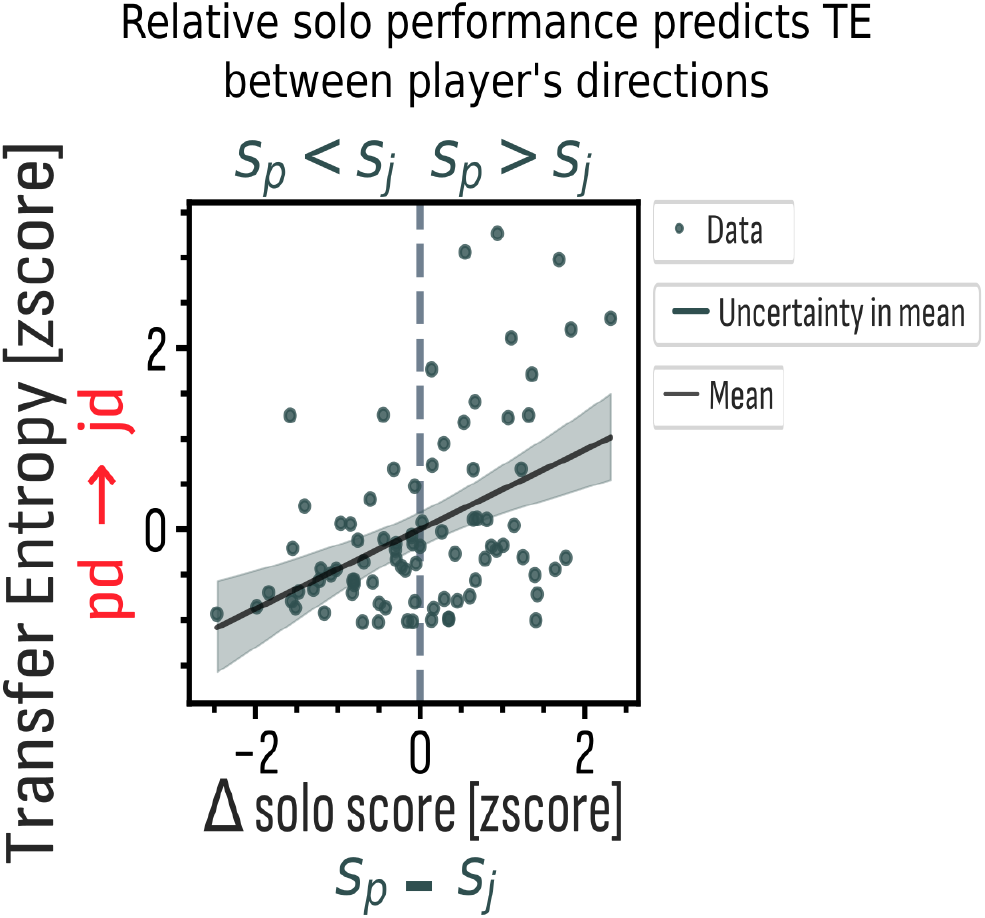
Posterior mean and 94% HDI for the regression of transfer entropy on Δ solo score. Transfer entropy from the partner’s joystick direction (pd) to the subject’s joystick direction (jd) is shown as a function of Δ solo score. Relative solo performance was defined as the difference between the partner’s solo score (*S*_*p*_) and the subject’s solo score (*S*_*j*_).

Taken together, these results indicate that subjects’ scores improve with greater social information obtained from their partner’s joystick direction and that this effect maybe based on subjects selectively acquiring more information from partners who outperform them.

### 4.3 Stimulus coherence modulates information transfer to player’s direction

Our previous results showed that subjects integrate both stimulus and social information sources in a behaviourally relevant manner, thereby improving their task performance (Fig. 4). Here, we investigated whether subjects’ reliance on their social partners is modulated by the reliability (i.e., noise level) of the stimulus.

Specifically, we asked whether the amount of information subjects obtained from their social partners is influenced by stimulus coherence, and whether this amount of information also differs between computer partners (i.e., in *H*_*C*_ dyads) and human partners (i.e., in *H*_*H*_ dyads). The latter question was motivated by previous work [5] reporting an improvement in human players’ accuracy and scores across all coherence levels when playing with a computer compared to a human partner, showing that the presence of a highly reliable “social” signal from a computer player could drive behavioral changes.

To investigate this, we estimated the amount of information transferred from the stimulus and partners’ direction to the players’ joystick direction (see section 3.10). We estimated the differences in TE between each pair of coherence levels for each link (i.e., sd → jd and pd → jd) using a Bayesian regression approach (see Fig. S3 and S4 for the forest plots showing the posterior distribution of the difference in TE between coherence levels).

As expected, increasing stimulus coherence was associated with greater transfer entropy from stimulus direction to players’ joystick direction (see *TE*_*sd*→*jd*_ in Fig. 6)

**Figure 6:**
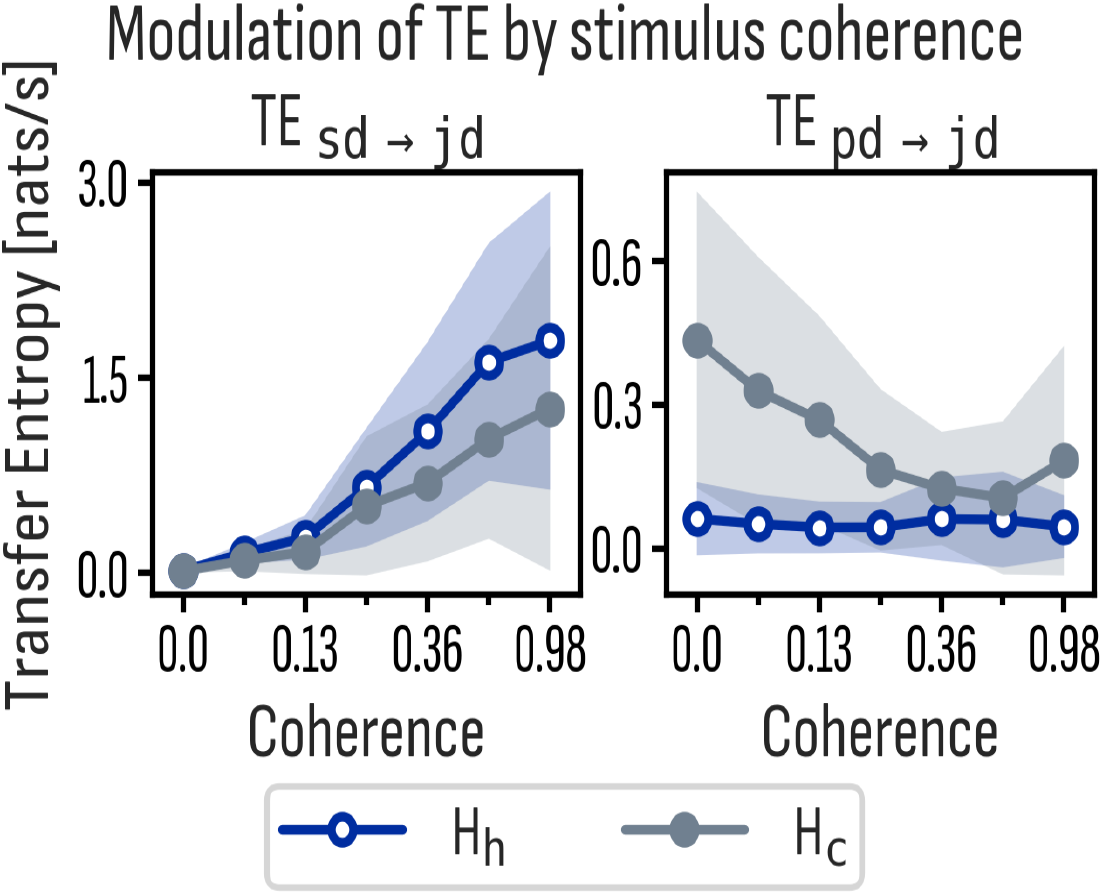
Stimulus coherence modulation of transfer entropy. Transfer entropy per coherence level across all subjects for stimulus direction to joystick direction (*sd* →*jd*) and partner’s direction to joystick direction (*pd* → *jd*) links. Color-coded for the type of dyad, i.e., human-human (*H*_*h*_) or human-computer (*H*_*c*_).

Conversely, as stimulus coherence decreased, we observed an increase of transfer entropy from the partners’ direction (see *TE*_*pd*→*jd*_ in Fig. 6). This latter effect was present only in the human-computer dyads, suggesting that subjects’ ability to rely on the “social” information under decreasing stimulus coherence (noisier stimulus) was only possible when the partners’ direction represented a highly reliable signal, as was the case with a computer player. Taken together, these results suggests that subjects can flexibly adapt from which sources they acquire information based on their reliability.

### 4.4 Subjects respond faster to a change in partners’ direction than to stimulus direction and partners’ tilt

We observed that subjects responded to a change in their partners’ direction faster than to both, a change in stimulus direction and in their partners’ tilt.

Specifically, subjects’ joystick direction responded to a change in partners’ direction 415 ms faster ([281, 552] ms 94% credible interval) than subjects’ joystick tilt to a change in partner’s tilt (Fig. 7). If we accept an interpretation of joystick tilt as a measure of perceptual confidence, this result suggests that subjects ability to respond to a change in their partners’ confidence requires a longer integration period than a response to their direction.

**Figure 7:**
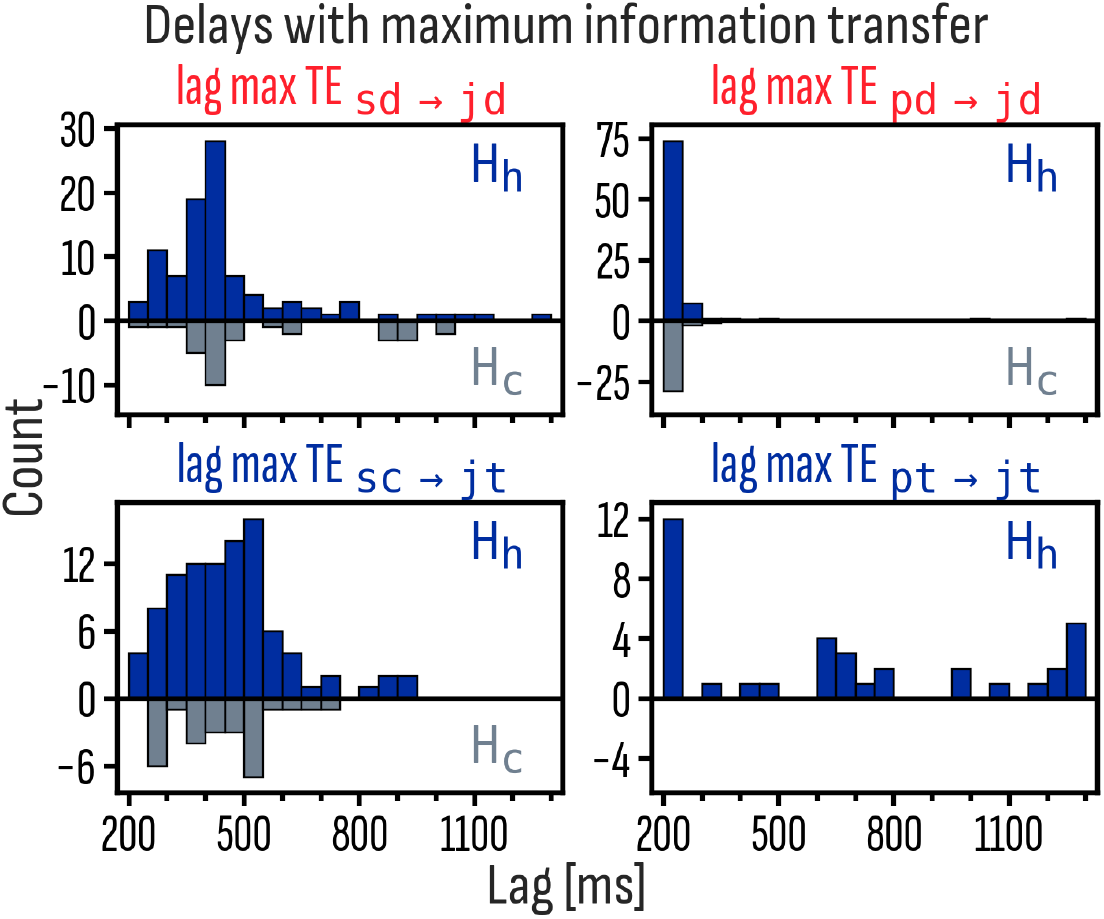
Distribution of lags with maximum information transfer. Lags with maximum transfer entropy for stimulus direction to joystick direction (*sd*→ *jd*), partner’s direction to joystick direction (*pd* →*jd*), stimulus coherence to joystick tilt (*sc* →*jt*) and partner’s tilt to joystick tilt (*pt*→ *jt*). Subjects response to a change in their partner’s direction was faster than to a change in partner’s tilt or stimulus direction.

Additionally, subjects’ joystick direction responded to a change in partners’ direction on average 239 ms ([196, 281] ms 94% credible interval) faster than to a change in stimulus direction (Fig. 7). Note that due to the use of a multivariate approach the respective information transfers from stimulus direction and partners’ joystick direction, as well as the lags of information transfer reported here, can be cleanly separated. These lags have to be interpreted as sequential lags, i.e. a subject first reacts to the change in stimulus direction with a certain lag, and the partner does so as well, then – once the partners’ change in response becomes visible – the subject reacts also to this second change, and the reaction time (as measured by the information transfer lag) is considerable shorter than to the initial reaction time to the stimulus change - possibly due to priming effects. The posterior distribution of the differences in lags between pairs of links is reported in Fig. S5 in the Supporting Information.

## 5 Discussion

As laid out in the introduction, research into social interactions that unfold continuously in time is an emergent field of research aimed at improving ecological validity. Yet, this field also requires novel methodological approaches to fully exploit the benefits of using continuous-time response behaviours in social settings instead of previous discrete, trial-based paradigms. Here, we combined a continuous perceptual decision making task in a dyadic setting [5] with information theory to study how human subjects continuously integrate partner information into their own behavior. Specifically, to understand how much information about a partner’s behaviour a subject integrates and at which latencies it does so, we used a multivariate transfer entropy analysis [11, 9, 22] which separates time-continuous stimulus- from time-continuous partner-based influences on a subject’s behaviour.

As a first result we found that real-time social signals predominantly induced interactions between corresponding behavioural dimensions, i.e. a partner’s direction assessment influenced the subjects choice of direction, while an indicator of the partner’s confidence influenced the subjects confidence, but there were only minimal cross-over influences between dimensions (i.e. the direction and the confidence dimensions).

Second, subjects integrated the above social information with information from the stimulus for their behavioral choices, as indicated by the fact that the actual amount of information transferred from both, social and stimulus sources, correlated with subjects’ task performance.

Third, subjects exhibited flexibility in gathering information from either stimulus or social sources depending on their relative reliability. In particular, when social information was highly reliable (e.g., when provided by a computer agent), subjects relied more heavily on social sources as the reliability of the stimulus decreased.

Finally, we observed that subjects’ reported direction and confidence were modulated by social information with distinct time delays. Specifically, subjects integrated their partners’ direction into their own joystick response 415 ms faster than they integrated their partners’ confidence into their confidence reports.

All of these results were obtained directly from reconstructing the network of interactions and their latencies using the model-free approach of multivariate transfer entropy and required no specific modeling choices. While we think that ultimately behavioral interactions must be carefully modeled to cast our understanding into a rigorous form, we consider the model-freeness of the information theoretic approach a distinct advantage to gain a first foothold into an understanding of continuous-time social interactions under more realistic settings.

### 5.1 Subjects integrate stimulus and social information in a behaviourally relevant manner in the CPR task

#### 5.1.1 Behaviorally relevant transfer of social information without instructions to interact

The CPR task used in our study critically differs from previous studies that investigated the behavioral consequences of accessing social information in dyadic settings; in the earlier studies described below subjects were explicitly instructed to interact, while in our task, payoffs were independent, and subjects were not required to communicate or coordinate. It is thus natural to ask whether both types of studies show similar outcomes.

For studies with instructed interaction it has previously been shown that partners influence task performance, for example through the communication of confidence [1, 2, 4]. Performance improvements have been observed when partners are equally expert [4], or when the higher-confidence partner’s choice is selected in contexts where participants tend to be correct [2]. By contrast, performance decreases have been reported when participants are prone to error [2], when expertise is unequal [4], or when group diversity is reduced without accuracy gains [23].

Our results agree with previous findings despite the fact that interactions were not instructed in our case. In particular, we demonstrated that subjects still integrated social and stimulus information in a behaviorally relevant way—that is, in a manner that influenced task performance. Specifically, the SSI model, which includes both stimulus and social transfer entropy predictors, outperformed the stimulus-only SI model in terms of out of sample prediction, indicating that social information contributed to predicting task performance.

Moreover, performance increased with greater information obtained from both, stimulus and social sources, suggesting that participants spontaneously integrated their partner’s behavior in a beneficial way even when interaction was not required.

Consistent with this interpretation, we found that higher acquisition of social information from a partner’s joystick direction was associated with higher subject scores, and that subjects paired with better-performing partners acquired more information from their partner. Accordingly, the amount of information acquired from the better performing partner increased with the difference in the skill levels between dyad members (as measured by their task performance in a solo setting).

#### 5.1.2 Classifying interaction types from the transfer entropy graph

In realistic examples of dyadic interactions there is a variety of social interactions that are often observed: mutually coordinated behaviour, leader-follower dynamics, or independently acting agents. As such a variance of behaviours is desirable for task designs for the research of social interactions, it is also desirable that the corresponding analysis methods readily detect it. In our novel analysis the graph of significant information flows in a dyad allows a straightforward differentiation of dyads into those with mutual influences between partners, leader-follower dynamics and uncoupled behaviours.

Our results also have connections with previous work using evolutionary simulations in a game-theoretic setting with varying visibility of a partner’s action. There, it was shown that the emergence of leader–follower dynamics indeed relied on the visibility of the others’ actions [24].

#### 5.1.3 Transfer of information on perceptual confidence

The communication of confidence between subjects has previously been reported during trial-based social interactions [1, 3, 4], and is also detected in our continuous report task by our information theoretic analysis.

In previous studies, human subjects were reported to align their confidence to those of their partners in group decision-making as an inexpensive strategy that requires just tracking others behaviour rather than inferring internal latent states [4]. Similarly, Pescetelli and colleagues [3] showed that regardless of subjects’ ability to report their internal uncertainty, highly confident individuals yielded the strongest social influence on dyadic outcomes. Similarly, we here observed that also real-time social interactions led to the transfer of information between subjects’ confidence indicators – in addition to the transfer of information between subjects’ choice of direction.

#### 5.1.4 Stimulus and social information sources are weighted differentially depending on their reliability

When playing against a highly reliable social partner (i.e., a computer player), subjects showed flexibility in relying on social information sources when the stimulus was highly noisy. We observed that as stimulus coherence decreased, information transfer from the stimulus direction decreased, while information transfer from the partner’s direction increased.

Humans’ ability to flexibly rely on different sensory information sources and weigh their contributions according to reliability has been extensively studied in the field of multisensory integration [25, 26, 27, 28].

For instance, when asked to discriminate the average value of a feature in an array of elements, participants were found to base their decisions primarily on evidence close to the array mean, while down-weighting or excluding evidence farther from the mean. This “robust averaging” strategy was interpreted as down-weighting less trustworthy evidence [25]. Similarly, weighting sensory cues according to their reciprocal variances has been reported during the integration of visual and haptic information [28].

Dynamic aspects of multisensory integration further show that humans are capable of flexibly adjusting their weighting of time-varying sensory inputs. Subjects have been found to reweigh modalities on a trial-by-trial basis depending on sensory reliability [26], and even to rapidly reweigh cues within a single trial [27].

In our task, subjects integrated two information sources from the same modality (i.e., visual), one of which carried social information. Consistent with subjects’ ability to flexibly weigh sensory input by reliability, we observed that subjects dynamically adapted over the course of a session, relying more on their partners as the stimulus became less reliable. This suggests that humans flexibly reweigh stimulus and social information according to the relative reliability of each source.

### 5.2 Confidence information requires longer integration time

Subjects’ choice direction responded faster than their confidence to changes in the corresponding partner’s dimension, suggesting that subjects required a longer accumulation period in order to incorporate, into their own behavioral reports, their partners’ confidence compared to their direction. Previous work has shown that while confidence judgments rely on the same sensory evidence as perceptual decisions, observers typically commit to perceptual decisions before trial termination, whereas confidence judgments are not prematurely terminated and instead continue accumulating evidence [29]. Moreover, distinct neural representations have been reported for perceptual decisions and confidence evaluations: perceptual decision representations cease incorporating additional evidence after commitment, whereas confidence representations continue to track evidence beyond decision commitment [30].

We also observed that subjects’ direction responded faster to changes in their partners’ direction than to changes in stimulus direction. This can be explained by the multivariate nature of our analysis and the sequential nature of two behavioral reactions. First subjects received stimulus information and integrate it into behavior. The information transfer from the partner is measured conditional on stimulus information and happens later in time – but then has a shorter latency. We interpret this effect as an adaptation of a subject’s initial perceptual decision, which is based on stimulus information, by information more rapidly extractable from their partners’ direction. The observed speedup is possibly due to behavioral priming as stimulus-direction changes and the partner direction-responses more frequently point to similar directions.

### 5.3 Considerations on the interplay of experimental design and information theoretic analyses

Information theoretic measures require the explicit or implicit estimation of (at least some properties of) probability distributions. This difficult task is further complicated if the relevant probability distributions change over time - as they are bound to do in trial-based designs with a fixed temporal structure. While there are ways to deal with such nonstationarities in subsequent statistical testing via suitable surrogate data (e.g. see [13] and references therein), continuous-time experimental designs with ongoing interactions and stimulus changes offer the possibility to obtain approximately stationary distributions, rendering the combination of continuous-time interactions and information-theoretic analysis beneficial in both directions.

## 6 Conclusion

We analyzed a continuous psychophysics task in which subjects reported the direction of motion of a RDP and their confidence in these choices in a dyadic setting with real-time social feedback. Using transfer entropy, we quantified the amount of information transferred from both stimulus and social information sources.

We found that real-time social feedback induced interactions within the same behavioural response dimensions, enabling the integration of stimulus and social information and the transfer of behaviourally relevant information. Moreover, subjects showed the flexibility to acquire more information from their partners when stimulus reliability was reduced. Finally, time lags at maximum information transfer revealed that subjects integrated their partners’ choice direction faster than their confidence.

Here, we demonstrated how information theory can contribute to the study of cognition under naturalistic conditions, such as those examined with continuous psychophysics paradigms [6, 7, 8], where perception unfolds continuously over time, and where complex networks of behavioral interactions may be difficult to model *ad hoc*. Thus, our multivariate, model-free information-theoretic approach may further advance the development of methods for studying naturalistic behaviour, and allow for an analysis of more complex designs [31].

## 7 Supporting information

### 7.1 Bayesian statistics

**Figure S1:**
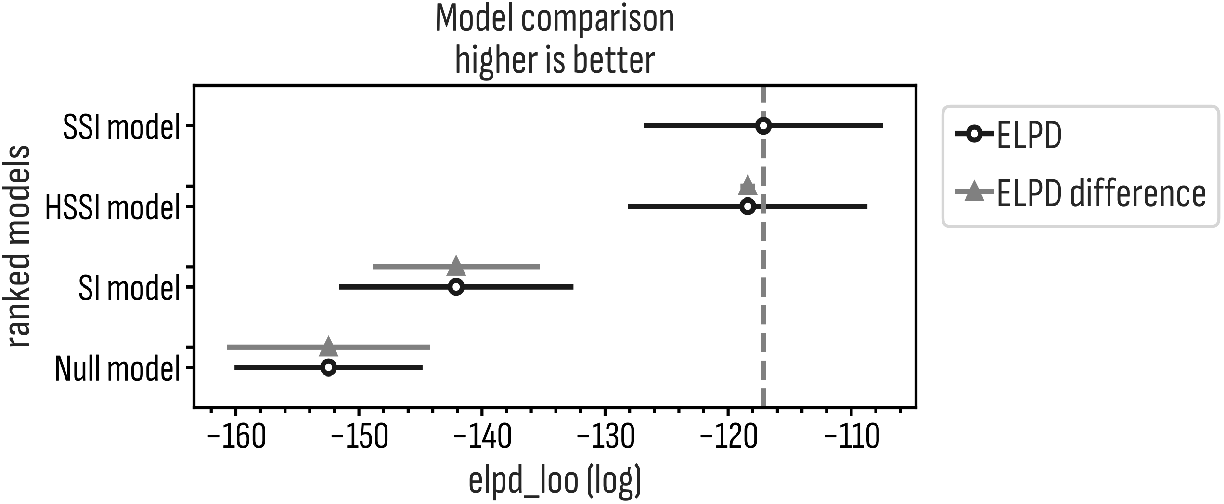
Model comparison using expected log pointwise predictive density (ELPD) estimated using leave-one-out cross-validation (loo). SSI and HSSI models perform equally well, with SSI slightly better, as indicated by the smaller ELPD. Models without social information (SI and Null) perform worse at predicting subjects’ score.

**Figure S2:**
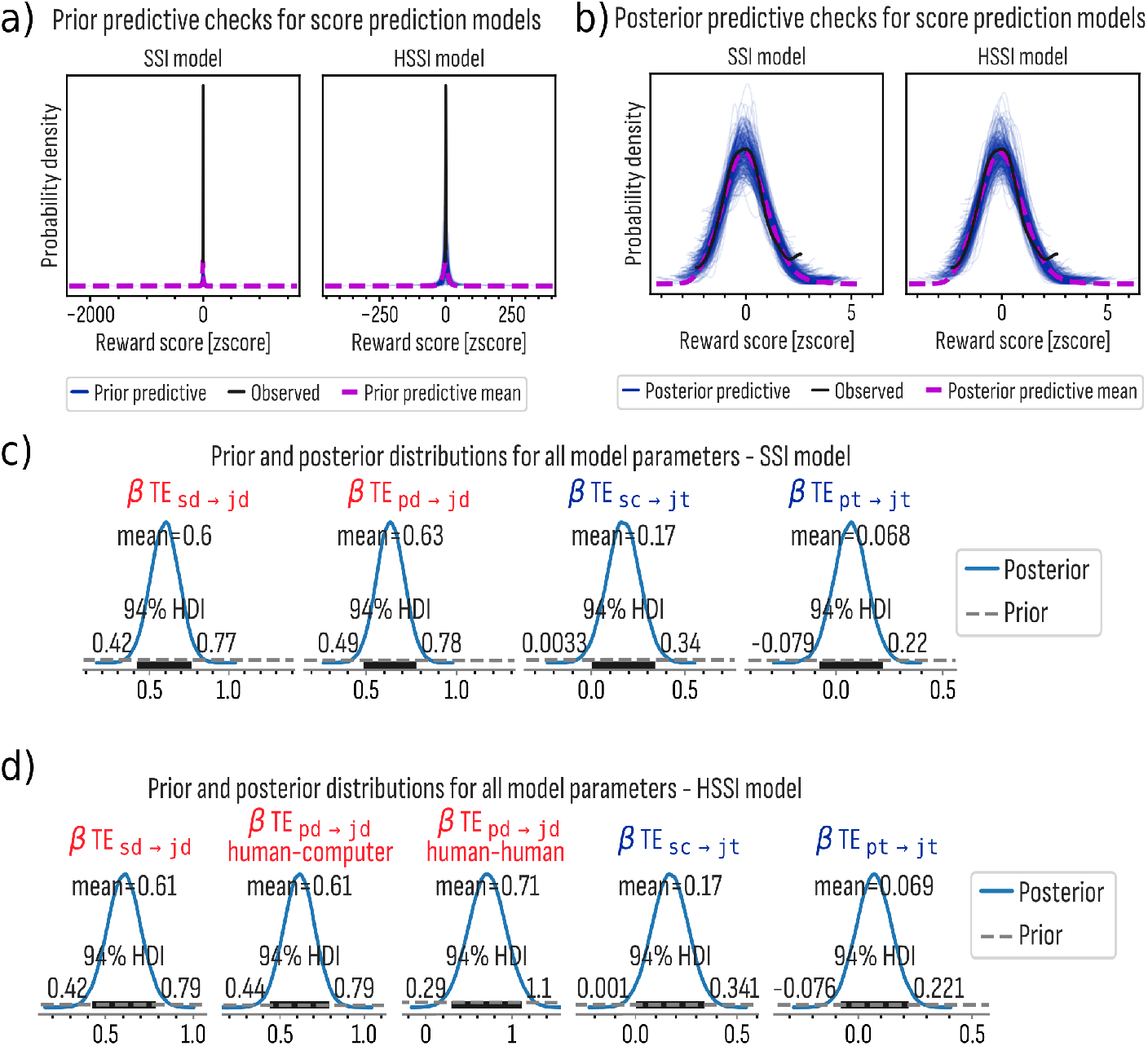
Prior and posterior predictive checks for SSI and HSSI models. (a) Prior predictive checks shows the range of plausible values for subjects’ scores as generated by the model before constraining it by the data. Prior values completely covers the subjects’ scores observed in the data. (b) Posterior predictive checks. After fitting the model to the data, subjects’ reward scores overlap with the observed values. (c-d) Prior (grey dashed) and posterior (blue solid lines) distributions for all model parameters for SSI (c) and HSSI (d) models. Priors were flat within the small range of values where the posterior distribution concentrates 94% of its values.

**Figure S3:**
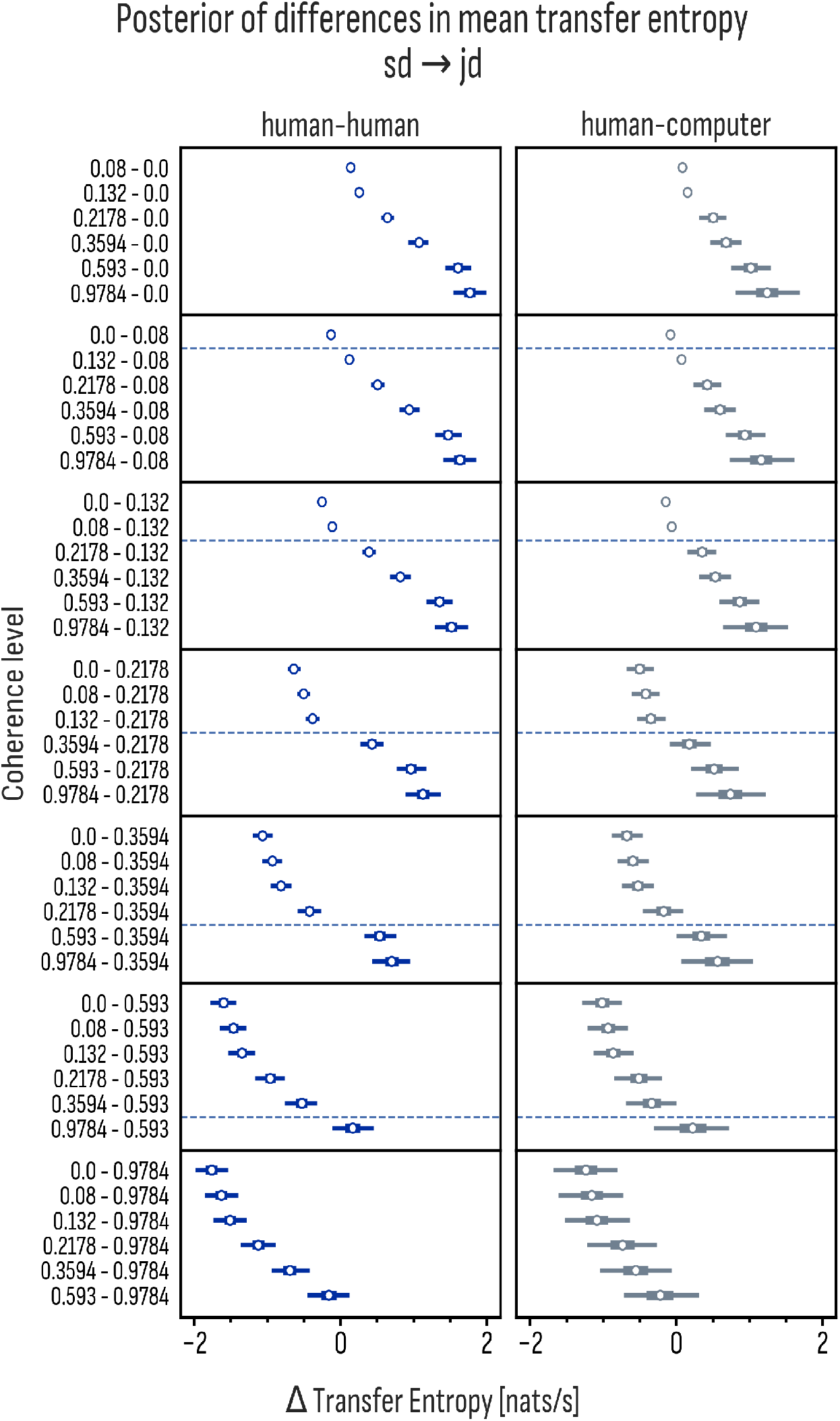
Posterior distributions of the difference in the mean TE for each pair of co-herence level for. *sd* → *jd* **link**. Transfer entropy increases with increasing stimulus coherence as indicated by the positive 94%HDI between two increasing coherence levels (e.g., 0.132 - 0.08). Both human-human and human-computer dyads presented the same effect.

**Figure S4:**
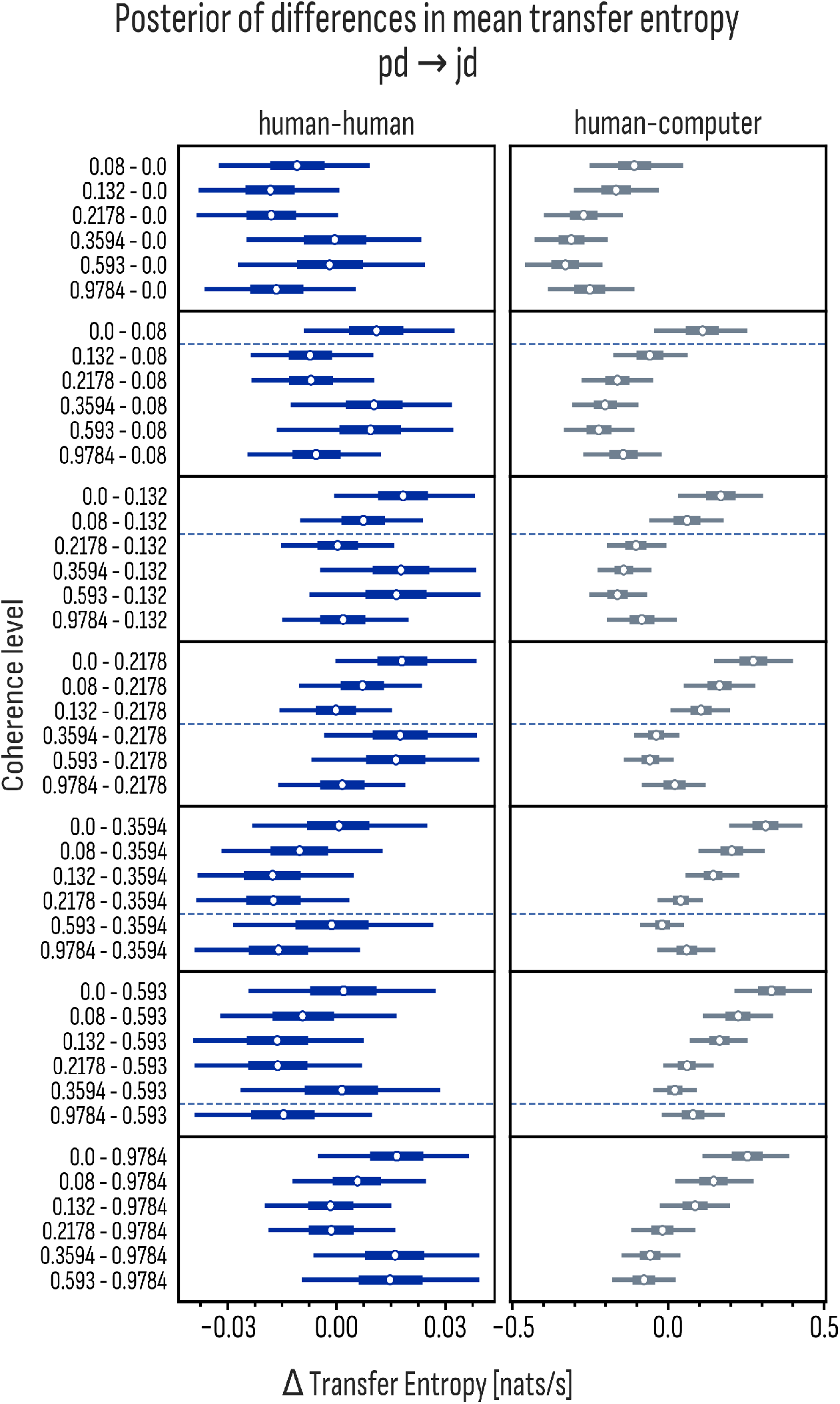
Posterior distributions of the difference in the mean TE for each pair of co-herence level for. *pd* → *jd* **link**. Transfer entropy decreases with increasing stimulus coherence as indicated by the negative 94%HDI between two increasing coherence levels (e.g., 0.132 - 0.08) for human-computer dyads. Stronger differences were observed between distant coherence levels (e.g., 0 - 0.978). No differences in TE were observed between coherence levels in the human-human dyads

**Figure S5:**
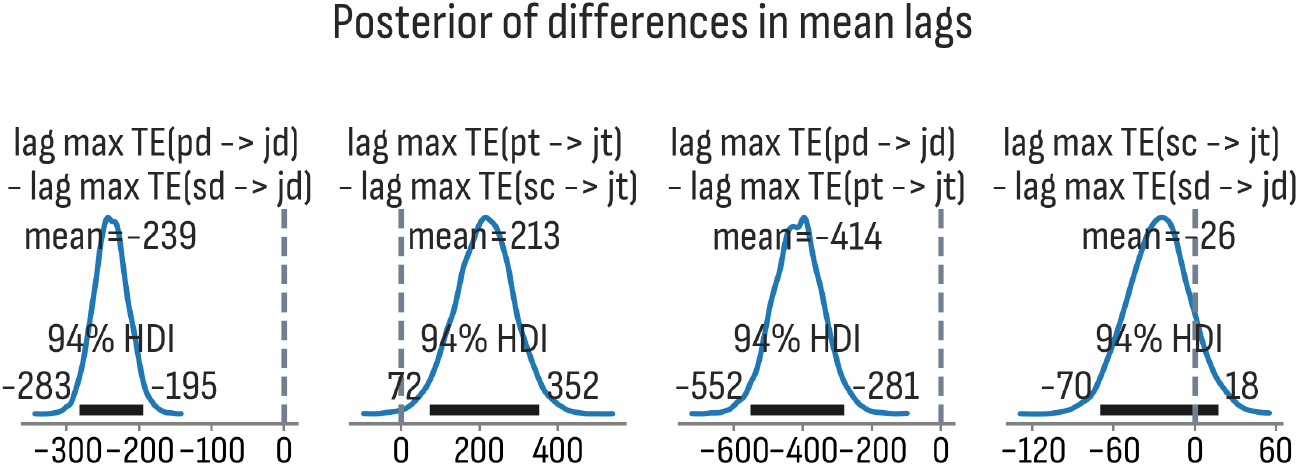
Posterior distribution of the difference in the mean lag for each pair of link. Faster lags with maximum information transfer were observed in the link pd → jd compared to sd → jd and pt → jt, as indicated by the negative 94%HDI. Smaller lags were observed in sc → jt compared to pt → jt, as indicated by the positive 94%HDI.

